# Chemosensory receptor copy-number evolution is not associated with host-plant specialization in butterflies

**DOI:** 10.64898/2026.11.05.762600

**Authors:** Sofia Danielson, Amna Ahmad, Zarley Rebholz, Kathy Darragh

## Abstract

Chemoreception plays an important role in how insects sense their environment and interact with other organisms. One important ecological role for chemosensory receptors is sensing chemical compounds to locate suitable host plants. Host-plant use is therefore expected to influence insect sensory system evolution as host-plant preferences changes and species become more generalized or specialized. Yet it remains unclear if species that are generalists require a larger chemosensory receptor repertoire. Here, we investigated the relationship between host-plant specialization and the evolution of olfactory receptor (OR) and gustatory receptor (GR) gene families across 87 butterfly species spanning six families. We annotated OR and GR repertoires and used phylogenetic comparative analyses to test whether receptor copy-number was associated with specialist versus generalist host-use strategies or with host-plant breadth. In addition, we reconstructed evolutionary transitions between generalist and specialist strategies and tested whether these transitions were associated with expansions or contractions in gene-family size. We found that OR and GR repertoire size did not differ between specialists and generalists, nor was it correlated with the number of host-plant families used by a species. We also did not find associations between gene family expansions or contractions and transitions between generalist and specialist strategies. Overall, we find that host-plant specialization is not associated with chemosensory receptor repertoire size across butterflies. We suggest that host-plant chemosensory adaptation may depend on other changes such as receptor tuning, expression differences, or changes in specific receptor lineages not captured by our analyses.

**Significance Statement:** Animal sensory systems are predicted to evolve in response to ecology, yet it is unclear whether insects that use a wider range of host plants require larger or more dynamic chemical-sensing systems. Across the genomes from 87 butterfly species, we found no association between host-plant breadth and either chemosensory receptor number or turnover. Our results suggest that host-plant adaptation is not explained by total receptor number alone and may instead depend on changes in particular receptors, their expression or tuning, and other ecological pressures shaping chemical sensing.

## Introduction

Chemosensory receptors play a critical role in facilitating how organisms perceive their environment. Insects possess several different chemosensory receptors; two of these, olfactory receptors (ORs) and gustatory receptors (GRs), allow the organism to detect and respond to chemical cues in their environments (Sokolinskaya et al. 2020). ORs primarily detect volatile organic compounds, whereas GRs generally detect non-volatile compounds through direct contact (Dahanukar et al. 2005). Repertoires of chemosensory receptors allow insects to discriminate amongst chemical stimuli and translate these signals into neural and behavioral responses. These repertoires play a key role in mediating behaviors important for reproduction and survival such as choosing mates (Smith and Breed 1995), finding host plants (Bruce et al. 2005), detecting prey or predators (Thomas et al. 2008), and defense mechanisms (Mondet et al. 2021).

Insect herbivores range from specialists that use a single or limited number of host plant species to generalists that feed or oviposit across a much wider range of plants (Ehrlich and Raven 1964). Host-plant recognition relies heavily on chemical cues, including both individual compounds and the composition and relative abundance of compounds within complex blends (Bruce et al. 2005). The chemical profiles of different plant taxa can vary considerably, influencing the diversity of chemical cues encountered by species during host-plant recognition. Therefore, specialists and generalists may encounter different ranges of plant chemicals in host assessment, which may be reflected in changes to their chemosensory receptor repertoires.

Host-plant specialization has been associated with changes in chemosensory repertoires, but findings are not consistent across insect lineages. Generalists may require more receptors as they are exposed to a broader range of plant compounds (Xu et al. 2016; Suzuki et al. 2018; Wang et al. 2022). Similarly, some specialists have been associated with reduced chemosensory repertoires, consistent with the idea that they would need to detect fewer plant compounds (Robertson and Wanner 2006; McBride and Arguello 2007; Ramasamy et al. 2016; Reisenman et al. 2023). However, some studies find no association between host breadth and receptor repertoire size (Gardiner et al. 2008; Yin et al. 2022; Toh et al. 2026). Many of these studies focus on single species or closely related taxa, leaving the relationship largely untested across broader phylogenetic scales.

Butterflies provide a strong comparative system for investigating the relationship between host-plant specialization and chemosensory receptor evolution. Interactions between butterflies and their host plants have long been thought to play an important role in the evolution and diversification of the group (Ehrlich & Raven 1964). A recent phylogenomic study provides a well-resolved phylogeny and natural history information on host-plant use (Kawahara et al. 2023), and extensive genomic resources in butterflies provide a framework to study the molecular bases of these interactions. Previous comparative studies in butterflies have found mixed evidence for a relationship between host-plant usage and chemosensory repertoires (Xu et al. 2016; Suzuki et al. 2018; Yin et al. 2022), making it unclear whether host specialization produces predictable changes in receptor repertoire evolution across the group.

Here, we investigate whether host-plant specialization is associated with the size of olfactory and gustatory receptor repertoires in butterflies. We annotated OR and GR genes in 87 species representing six butterfly families and used phylogenetic comparative analyses to examine the relationship between host breadth and receptor repertoire size. In addition, we investigated patterns of receptor gene-family evolution to determine whether host-plant specialization was associated with changes in OR or GR evolutionary dynamics. By expanding taxonomic sampling, our study aims to shed light on whether host-plant specialization is associated with chemosensory repertoire size and evolutionary dynamics across butterflies.

## Results

### OR and GR repertoire sizes vary across butterfly species

To identify ORs and GRs, we used a reference dataset of published sequences to search our newly generated genome annotations for each of our 87 target species. After filtering sequences for size, protein domain completeness, and 5-8 transmembrane domains, our final dataset contained 4274 ORs and 4115 GRs across 52 generalists and 35 specialist butterfly species (Fig. 1). GR repertoire size ranged from 20 in *Leptidea sinapis* to 118 in *Mechanitis polymnia* (mean ± SD, 47±21). OR repertoires showed less variation, from 32 in *Lamproptera curius* to 66 in *Eueides isabella* (mean ± SD, 49±7). Although mean repertoire sizes were similar between the two receptor families, GR repertoire size was more variable across species (Paired Pitman-Morgan test, t_85_ = 12.74, p<0.001).

**Figure 1.**
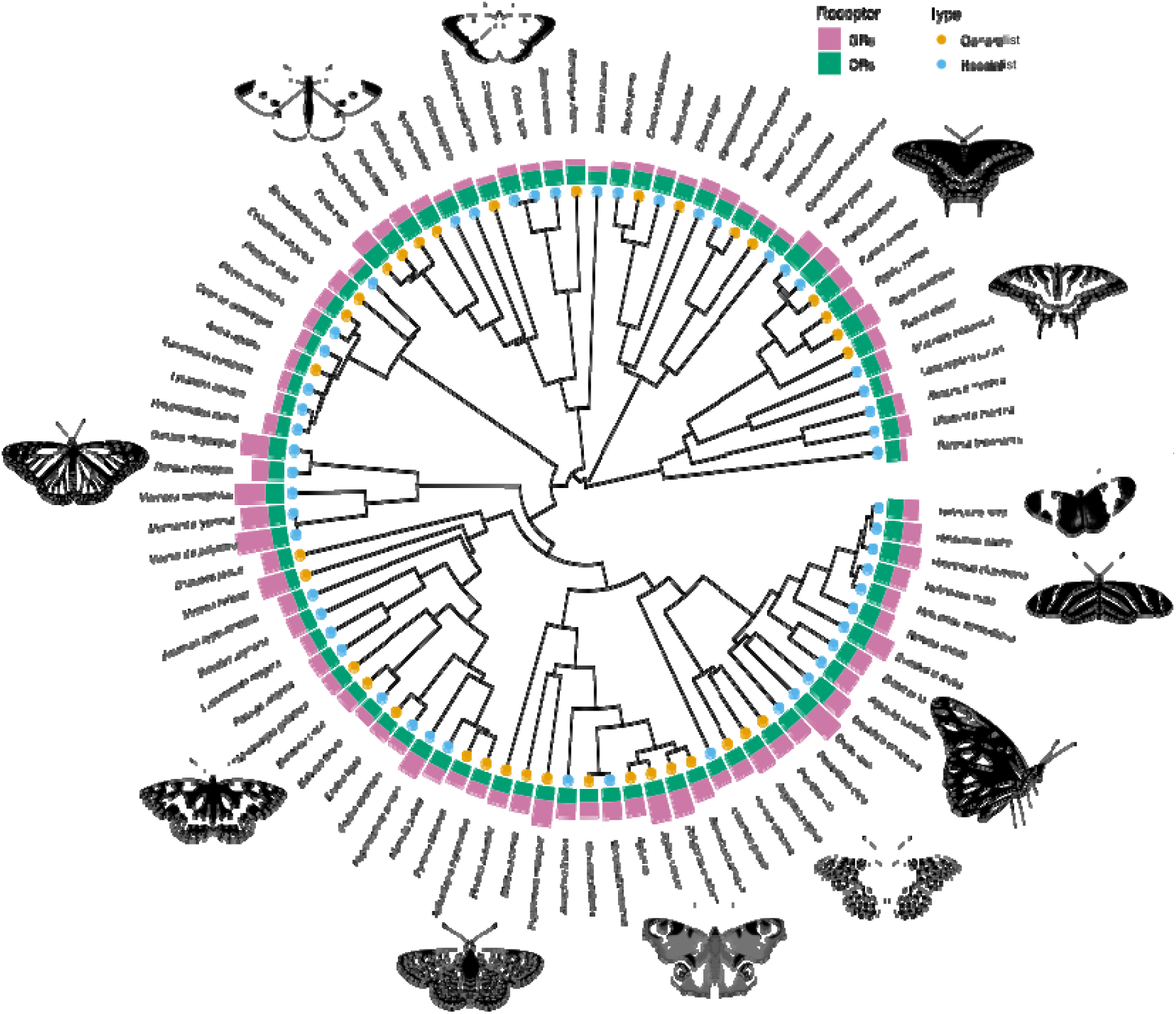
Phylogenetic distribution of olfactory and gustatory receptor repertoires across 87 butterfly species. The phylogeny includes 52 host-plant generalists and 35 specialists and was derived from Kawahara et al. (2023). Phylogeny tips indicate host-plant specialization category, and outer bars represent the number of gustatory receptors (GRs) and olfactory receptors (ORs) identified in each species.

We combined our sequences with previously published sequences from *Manduca sexta* to generate a gene tree for each receptor family (Figs. S1, S2). While *Manduca sexta* sequences were trimmed for downstream gene family analyses, they allowed us to identify previously characterized clades of receptors. For GRs, we identified CO₂, sugar, and fructose clades based on orthology with *Manduca sexta* sequences. For the ORs, we were able to identify the conserved ORCO clade and a pheromone-receptor clade within the broader phylogeny.

### Receptor repertoire size is not associated with host-plant specialization

We analyzed differences in receptor copy numbers between species that differ in host-plant specialization strategies, with generalist and specialist first treated as binary categories. We accounted for phylogenetic relationships using a previously published butterfly phylogeny (Kawahara et al. 2023). We found that GR and OR repertoire sizes were similar between generalists and specialists. Generalists had a mean of 47 GRs, compared with 47.5 in specialists, with repertoire size not differing significantly between specialization strategies (Fig. 2A, phylogenetic ANOVA, F*_1,85_* = 0.014, P= 0.905). Similarly, generalists had a mean of 48.5 ORs, compared to 49.5 in specialists, with no significant difference between groups (Fig. 2B, phylogenetic ANOVA, F*_1,85_* = 0.357, P= 0.552).

**Figure 2.**
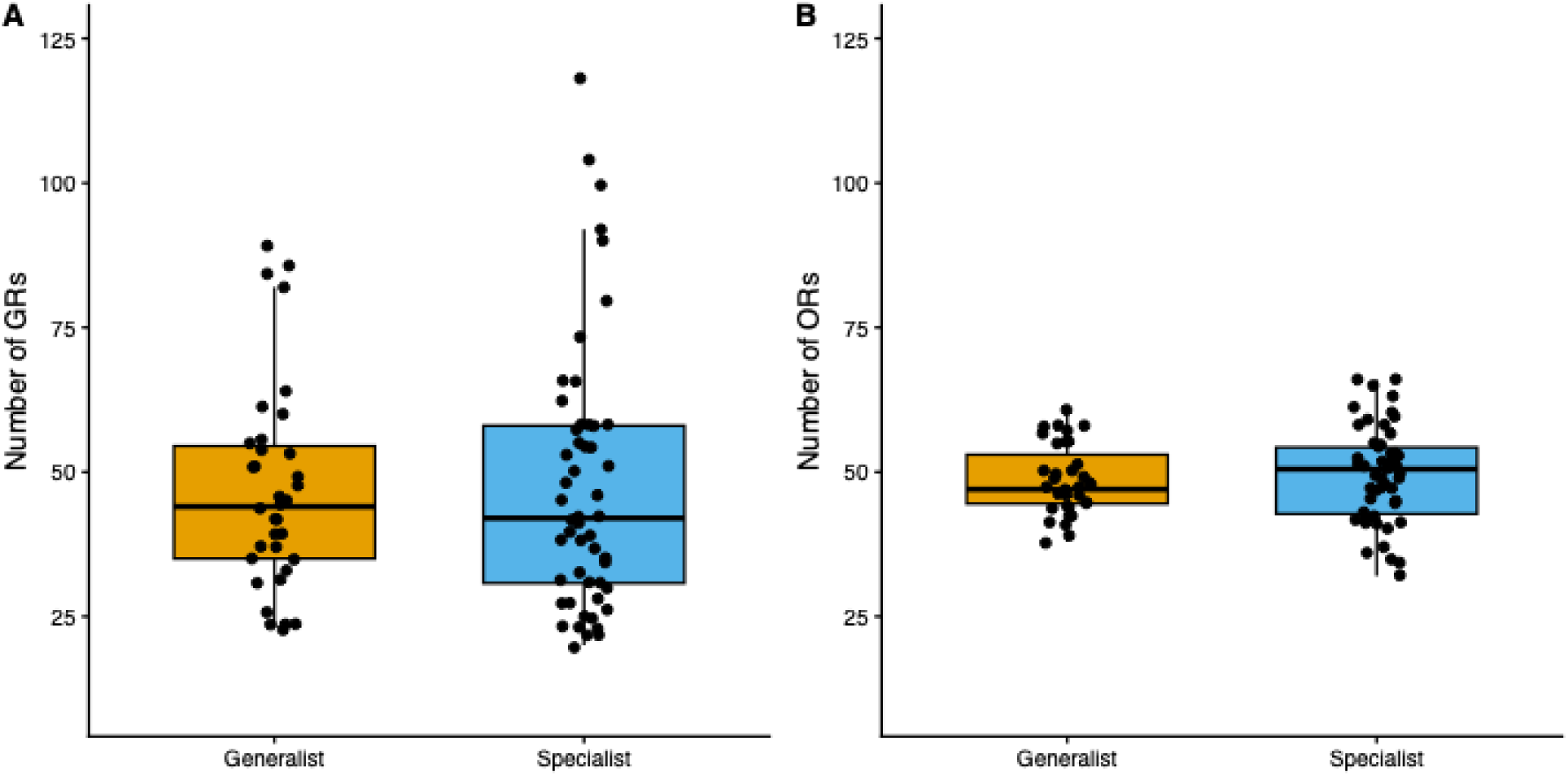
Olfactory and gustatory receptor repertoire sizes in host-plant generalist and specialist butterflies. (A) Gustatory receptor (GR) repertoire size did not differ significantly between generalists and specialists (phylogenetic ANOVA, *F_1,85_* = 0.014, *P* = 0.905). (B) Olfactory receptor (OR) repertoire size also did not differ significantly between specialization groups (phylogenetic ANOVA, *F_1,85_* = 0.357, *P* = 0.552).

Next, we treated host breadth as a continuous variable, using the number of host-plant families for each species as a measure of host breadth (Kawahara et al. 2023). After accounting for phylogenetic effects using phylogenetic independent contrasts (PICs), host breadth was not significantly correlated with GR repertoire size (Fig. 3A, Spearman’s correlation, ρ = 0.057, *P* = 0.601), nor OR repertoire size (Fig. 3B, Spearman’s correlation, ρ = −0.151, *P* = 0.163). Overall, we detected no association between host-plant specialization and chemosensory repertoire size using either binary specialization categories or a continuous measure of host breadth.

**Figure 3.**
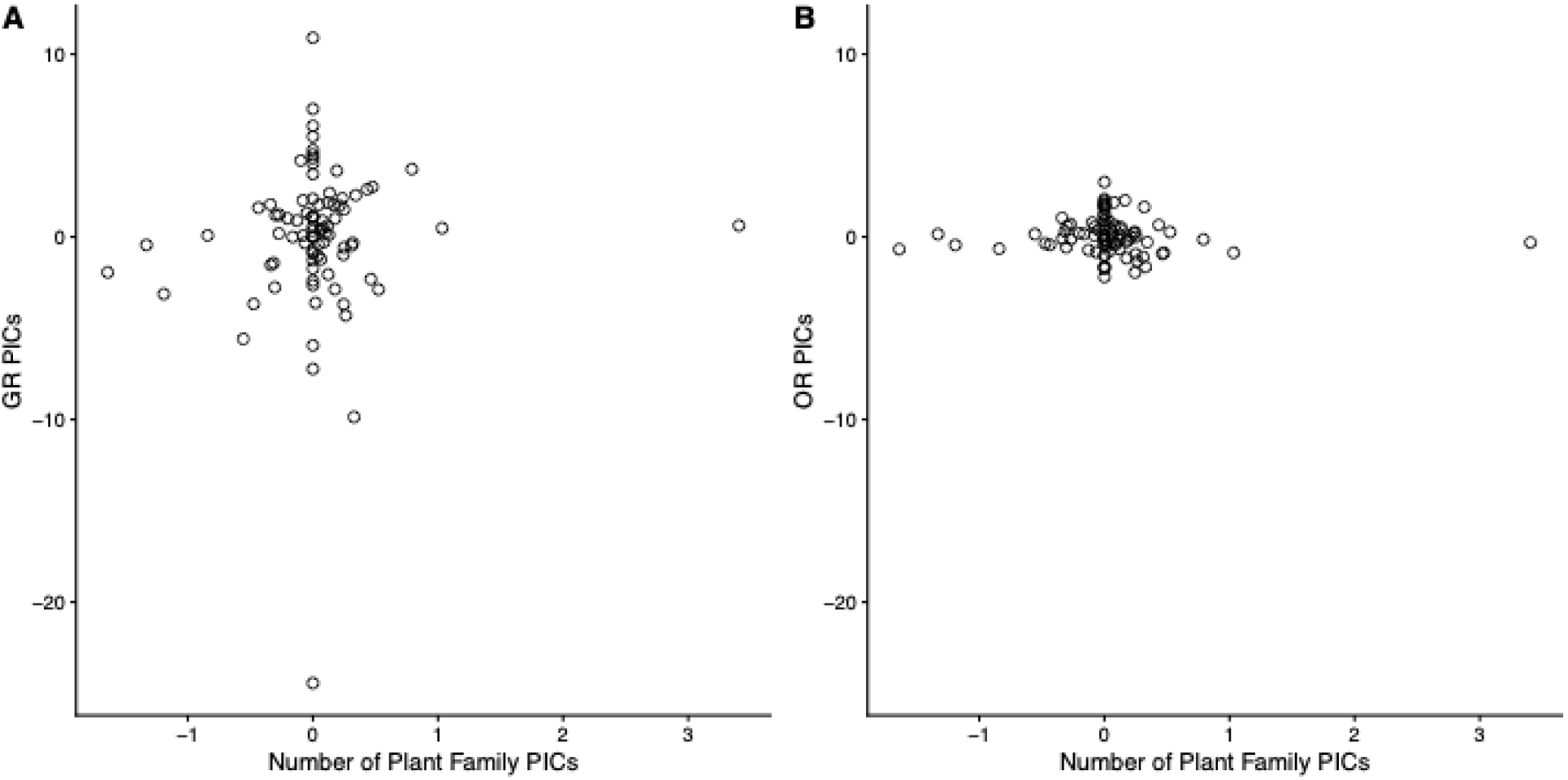
Relationship between host-plant breadth and chemosensory receptor repertoire size. (A) GR repertoire size was not significantly correlated with host breadth (Spearman’s ρ = 0.057, *P* = 0.601). (B) OR repertoire size was not significantly correlated with host breadth (Spearman’s ρ = −0.151, *P* = 0.163). PICs, phylogenetically independent contrasts.

### Changes in receptor repertoire size are not associated with transitions in host-plant specialization

We next tested whether evolutionary transitions in host-plant specialization were associated with changes in OR or GR repertoire size. To do this, we used ancestral state reconstruction of host-plant specialization to identify branches where transitions occurred and CAFE to infer expansions and contractions in OR and GR gene families along the species tree (Mendes et al. 2020). We combined these datasets to ask whether transitions from generalist to specialist or specialist to generalist are associated with expansions or contractions in GR and OR families.

We identified 17 transitions from specialist to generalist and nine transitions from generalist to specialist across the phylogeny (Fig. S3). CAFE-inferred expansions and contractions in OR and GR repertoires associated with these transitions are shown across the complete phylogeny in Fig. S4. We found no significant association between the direction of host-plant specialization transitions and changes in chemosensory receptor repertoire size (Fig. 4). Net changes in GR repertoire size did not differ significantly between transitions (ANOVA, *F*₁,₂₄ = 1.41, *P* = 0.247; Fig. 4B). OR repertoire changes also did not differ significantly between transition directions (*F*₁,₂₄ = 3.39, *P* = 0.078; Fig. 4C). Although the exact branch-level reconstructions differed, we obtained consistent results using gene-tree/species-tree reconciliation with reconcILS (Mishra et al. 2024). Net changes in repertoire did not differ between transition directions for either GRs (*F*₁,₂₄ = 0.019, *P* = 0.891) or ORs (*F*₁,₂₄ = 0.330, *P* = 0.571; Fig. S5). Thus, despite differences in the specific gene-family changes inferred by the two approaches, neither analysis provided evidence that transitions in host-plant specialization were associated with changes in GR or OR repertoire size.

**Figure 4.**
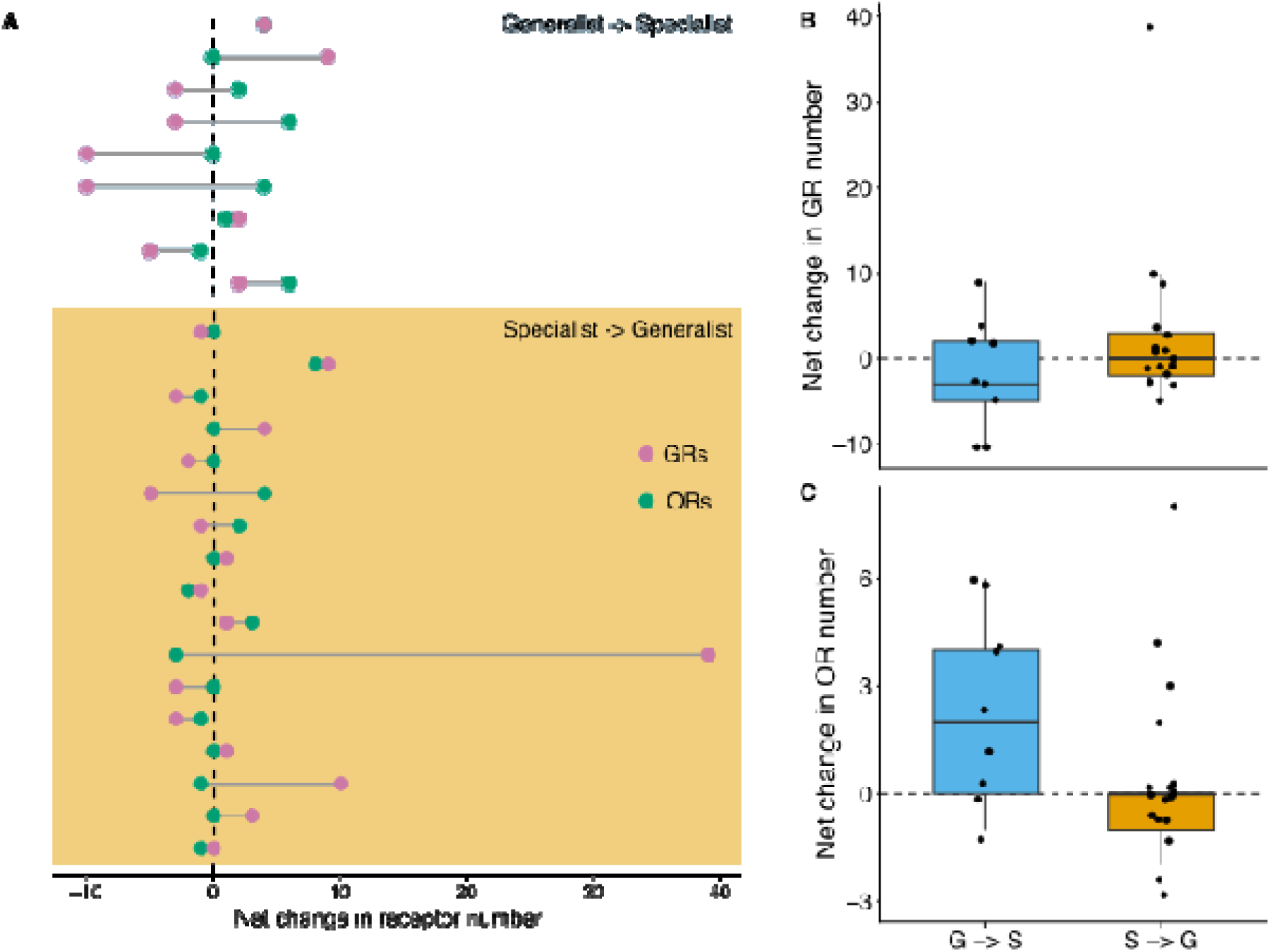
Changes in chemosensory receptor repertoire size associated with transitions in host-plant specialization. (A) Net changes in olfactory receptor (OR) and gustatory receptor (GR) gene-family size inferred by CAFE for branches undergoing transitions from generalist to specialist (blue) or specialist to generalist (orange). Each row represents an independent host-specialization transition; points indicate net OR and GR change, and connecting lines identify changes occurring on the same branch. Positive values indicate net expansion, and negative values indicate net contraction. (B–C) Distribution of net changes for G→S and S→G transitions for GRs (B) and ORs (C), respectively. Net changes did not differ significantly between transition directions for either GRs (ANOVA, *F*₁,₂₄ = 1.41, *P* = 0.247) or ORs (*F*₁,₂₄ = 3.39, *P* = 0.078). Points represent individual transition branches, and dashed lines indicate no net change.

## Discussion

In this study, we found that host plant specialization is not associated with olfactory and gustatory receptor copy-numbers in butterflies. Not only was repertoire size not different between specialists and generalists, but it was also not significantly correlated with host-plant breadth. Furthermore, net changes in OR and GR copy-number did not differ significantly between transitions toward specialization and transitions toward generalism. Previous comparative studies within genera or within tribes also did not find an association between chemosensory receptors and host plant specialization (Yin et al. 2022; Toh et al. 2026). Our results extend these findings across a much broader phylogenetic sampling of butterflies, encompassing six families. These results suggest that host-plant specialization does not predict total chemosensory receptor repertoire size or copy-number changes across butterflies.

Although not associated with host-plant specialization, we found that GRs exhibit higher variation in gene copy-number in the species we sampled, compared to ORs. This may reflect differences in the selective regimes or functional constraints of the two families. This finding is consistent with previous comparative work in Lepidoptera which suggested that there are more frequent expansions in GRs than ORs (Engsontia et al. 2014). Similarly, higher turnover of GRs than ORs has been described in *Heliconius*, including extensive GR duplication and greater copy-number variation in duplicated GR lineages (Briscoe et al. 2013). Comparative studies of *Drosophila* chemosensory receptors have similarly reported higher gene turnover and weaker selective constraint in GRs than ORs (Gardiner et al. 2008; Sánchez-Gracia et al. 2009). These findings suggest that GR repertoires may be more evolutionarily labile than OR repertoires, although the mechanisms underlying the greater variation observed in our dataset remain unclear.

The lack of association between host plant specialization and size of GR and OR repertoires does not mean that changes in gene family size are not important for host-plant adaptation. It is possible that changes in copy-number could be concentrated in specific GR and OR clades rather than repertoires as a whole. For example, the generalist *Vanessa cardui* exhibits higher rates of gene duplication in specific GR subfamilies but not in the conserved sugar and CO₂ receptor clades (Suzuki et al. 2018). In *Heliconius melpomene*, recently duplicated GRs are disproportionately expressed in female legs, suggesting that these lineage-specific duplications may play a role in host-plant assessment (Briscoe et al. 2013). Alternatively, factors unrelated to copy-number could be involved in chemosensory adaptation accompanying shifts in specialization.

Coding changes affecting receptor tuning and sensitivity are one mechanism by which host adaptation could occur without changes in repertoire sizes. Host-plant specialization behavior in *Drosophila sechellia* can be partly attributed to changes in the tuning of their Or22a receptor which results in high sensitivity to methyl esters (Auer et al. 2020). Similar patterns have been found in lepidopterans where two generalist species, *Spodoptera littoralis* and *S. litura*, and one specialist species, *S. picta,* demonstrate conservation of receptor repertoire despite having differing levels of host-plant specialization. Despite this conservation, there were notable differences in ligand specificity between species, with *S. picta’s* OR29 receptor exhibiting a heightened sensitivity to compounds emitted by their respective host plant (Comte et al. 2026). These studies demonstrate that receptor tuning can play a key role in host-plant specialization without changes in chemosensory receptor repertoires.

Additionally, regulatory changes could also facilitate host-plant adaptation by affecting receptor expression levels. For instance, the specialist *Drosophila sechellia* exhibits more rapid evolution of chemosensory gene expression and upregulation of multiple chemosensory genes when compared to closely related generalist lineages (Kopp et al. 2008; Shiao et al. 2015). While not specifically related to differences in host breadth, changes in receptor expression have also been associated with host shifts in *Drosophila* and leaf beetle species (Crowley-Gall et al. 2016; Wang et al. 2018) More generally, species with a similar GR and OR repertoire can show substantial differences in chemosensory receptor expression (Wang et al. 2023). Thus, host-plant adaptation can involve changes in gene expression without requiring accompanying changes in repertoire size.

Chemosensory receptor repertoires may also respond to ecological pressures unrelated to host-plant preferences. Adult butterflies use chemical cues for other tasks such as finding adult food sources using floral scent (Andersson and Dobson 2003), or finding and choosing mates using pheromones (Ehlers and Schulz 2023). Furthermore, the chemical background against which they are searching could also vary with time and habitat. For example, plant volatile emissions are released in higher amounts with more diverse blends during the day than at night (Lin et al. 2025), potentially exposing day-flying insects to different sensory environments. Different habitats are also likely to provide different sensory challenges, as plant community richness can impact the number and composition of volatiles in the environment (Kigathi et al. 2019; Ruggeri et al. 2026 June 21). Thus, chemosensory receptor evolution is likely influenced by more ecological factors than host-plant interactions alone.

Defining host-plant specialization by the number of families in which host plants occur is a relatively course proxy, and a limitation of our study. Taxonomic relatedness in plants does not necessarily correlate with chemical similarity. For example, there are plant families composed of species that are highly chemically divergent (Endara et al. 2015; Volf et al. 2015), and distantly related plants can share similar chemical profiles (Pichersky and Lewinsohn 2011). Future comparative analyses that incorporate host-plant chemistry rather than taxonomic similarity may reveal relationships between host-plant chemistry and chemosensory receptor evolution that we cannot detect in our current study.

Recent increases in genome availability allow us to study the evolution of chemosensory receptor gene families across rapidly expanding collections of species. These comparisons can reveal patterns of gene family evolution, but gene-family analysis alone cannot show how functional and regulatory changes are involved in host-plant adaptation. Because relatively few chemosensory receptors have been functionally characterized, our ability to interpret these analyses is limited. Experimental characterization of chemosensory receptors in more species and lineages will help us link gene-family evolution to receptor sensitivity, ecology, and function. Future studies integrating comparative genomics with receptor function, expression, and ecological data will help clarify the role of chemosensory receptor evolution in host-plant adaptation.

## Materials and Methods

### Gene family annotation

Genome assemblies for 87 species were obtained from NCBI and annotated using Helixer with the invertebrate model (Stiehler et al. 2021). To identify ORs and GRs, we compiled a reference dataset of published lepidopteran ORs and GRs from the following species: *Bombyx mori* (Wanner and Robertson 2008; Tanaka et al. 2009), *Heliconius melpomene* (Briscoe et al. 2013; Toh et al. 2026), *Papilio machaon* (Yin et al. 2022), *Pieris brassicae* (Wang et al. 2023), and *Manduca sexta* (Koenig et al. 2015). Our reference dataset was then used to generate alignment files for each genome from Exonerate (Slater and Birney 2005). We then used InsectOR (Karpe et al. 2021) to predict ORs and GRs for each genome. We identified transmembrane domains using DeepTMHMM (Hallgren et al. 2022). Our final dataset included protein sequences of 300-500 amino acids and 5-8 transmembrane domains. For species missing ORCO in the final dataset, we manually annotated ORCO using Exonerate (Slater and Birney 2005). Variance in repertoire size was tested using a Pitman-Morgan test using the function var.test in the R package *PairedData* (Champely 2018).

### Gene family phylogenetic analyses

Sequences from each gene family were combined separately and aligned using MAFFT v7.508 (-maxiterate 1000, L-INS-I algorithm)(Katoh and Standley 2013). Genes from *Manduca sexta* were included to identify previously described clades but were otherwise trimmed out of gene trees for downstream analyses. Alignments were trimmed using trimAl (-gt 0.5 for GRs, 0.2 for ORs)(Capella-Gutierrez et al. 2009). Phylogenetic trees were constructed using IQ-TREE with the ModelFinder function to determine the best-fit model (Minh et al. 2013; Nguyen et al. 2015; Kalyaanamoorthy et al. 2017; Hoang et al. 2018). The OR phylogeny was rooted using the ORCO clade, and the GR phylogeny was mid-rooted. To visualize the gene trees, we used *ape, ggplot2, ggtree,* and *tidyverse* (Wickham 2009; Yu et al. 2017; Paradis and Schliep 2019)

### Species phylogeny and host-plant specialization

The species phylogeny and host-plant dataset were obtained from a previously published dataset (Kawahara et al. 2023). The phylogeny was pruned using the drop.tip function in the R package *ape* (Paradis and Schliep 2019). Butterfly species were classified as host plant generalists or specialists following (Kawahara et al. 2023) where 1 = generalist and 0 = specialist, with number of host plant families also obtained as a more continuous measure of host breadth (Kawahara et al. 2023). To visualize the number of ORs and GRs along with host-plant specialization data on the species tree, we used *ape, ggplot2, ggtree, ggtreeExtra,* and *tidyverse* (Wickham 2009; Yu et al. 2017; Paradis and Schliep 2019; Xu et al. 2021).

### Statistical analyses of receptor repertoire size

To test for a correlation between host-plant specialization and receptor number, we ran a phylogenetic ANOVA using the function aov.phylo in the *geiger* package (Pennell et al. 2014). We visualized the distributions of receptor repertoire sizes in generalists and specialists using boxplots. We additionally tested whether receptor numbers were associated with host-plant breadth as measured by the number of plant families used by each species. To account for non-independence among species, phylogenetic independent contrasts (PICs) were calculated for receptor repertoire size and number of host-plant families using the pic function in the *ape* package (Paradis and Schliep 2019). A Spearman’s rank correlation test was then used to test relationships between the PICs of host breadth and OR or GR repertoire size using the cor.test function in R.

### Gene family analyses

To test whether evolutionary transitions in host-plant specialization were associated with changes in chemosensory receptor repertoire size, we reconstructed ancestral specialization states across the species phylogeny. Ancestral host plant states were estimated using the ace function in the *ape* package (Paradis and Schliep 2019) under an equal-rates (ER) discrete-character model. Internal nodes were then assigned as specialist or generalist according to the state with the highest reconstructed likelihood.

Changes in OR and GR gene-family size were computed using Computational Analysis of gene Family Evolution (CAFE, version 5) (Mendes et al. 2020) which models gene-family evolution using a birth–death model across a species phylogeny. These changes in gene family size were mapped to the species phylogeny to associate the transition branches in host-plant specialization with gene family expansions and contractions. We tested whether net repertoire change differed between transitions from generalist to specialist and specialist to generalist for both ORs and GRs separately using ANOVAs. For visualization, transition branches were highlighted on the phylogeny and OR and GR changes plotted as bars above (showing expansions) and below (showing contractions). To improve visualization, square-root scaling was used.

An alternative approach to investigating the dynamics of gene family evolution is through reconciliation of the gene-tree and species-tree. To do this, we used reconcILS (Mishra et al. 2024). For each branch, we calculated net change in receptor repertoire size as the number of inferred duplications minus the number of inferred losses. reconcILS-inferred changes were mapped to the same branches used in the ancestral-state reconstruction, allowing us to identify changes associated with transitions in host-plant specialization. As with the CAFE analysis, we tested whether net OR and GR repertoire changes differed between transition directions using separate ANOVAs.

Multi-panel figures were created using the package *cowplot* (Wilke 2020). All statistical analyses were conducted in R version 4.6.1 (R Core Team 2026).

## Data Availability Statement

All data and code used in this study are available on Zenodo (10.5281/zenodo.23194266).

## Acknowledgements

We would like to thank Matthew Hahn and Sarthak Mishra for helpful discussions about the gene family analyses.

## Notes

### Competing Interest Statement

The authors have declared no competing interest.

