## Supplementary file for "Chemosensory receptor copy-number evolution is not associated with host-plant specialization in butterflies"

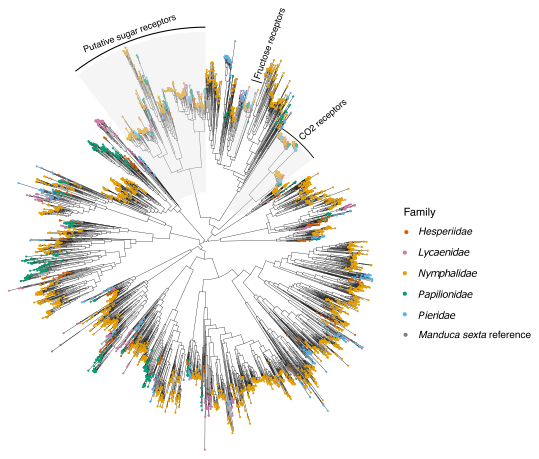
**Figure S1. Phylogeny of gustatory receptors (GRs) across 87 butterfly species.** The phylogeny was constructed in IQ-TREE using 4157 amino acid sequences with the LG model. Tip colors indicate butterfly family. Manduca sexta reference sequences were included to identify previously characterized receptor groups. Highlighted regions indicate the CO₂ receptor, putative sugar receptor, and fructose receptor clades. The phylogeny was midpoint rooted.

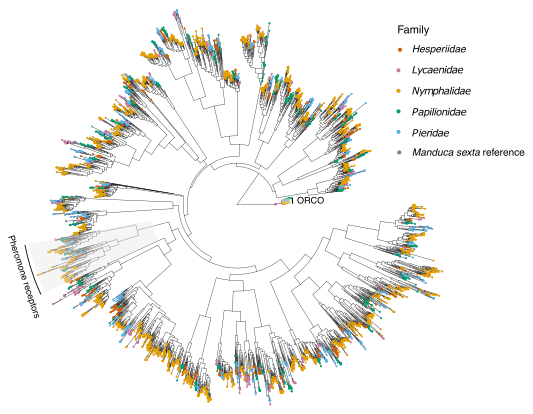

**Figure S2. Phylogeny of olfactory receptors (ORs) across 87 butterfly species.** The phylogeny was constructed in IQ-TREE using 4356 amino acid sequences with the LG model. Tip colors indicate butterfly family. Manduca sexta reference sequences were included to identify previously characterized receptor groups. Highlighted regions indicate the Orco and putative pheromone-receptor clades. The phylogeny was rooted using the Orco clade.

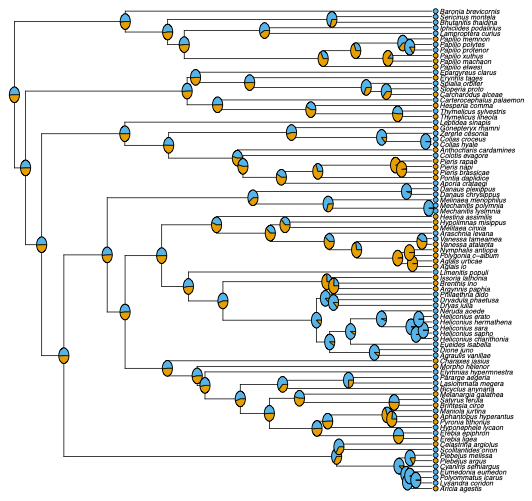
**Figure S3. Ancestral state reconstruction of host plant specialization across the butterfly phylogeny.** Tip states indicate observed host-plant specialization, with generalists shown in orange and specialists in blue. Pie charts at internal nodes represent the relative likelihoods of reconstructed generalist and specialist states under an equal-rates (ER) model.

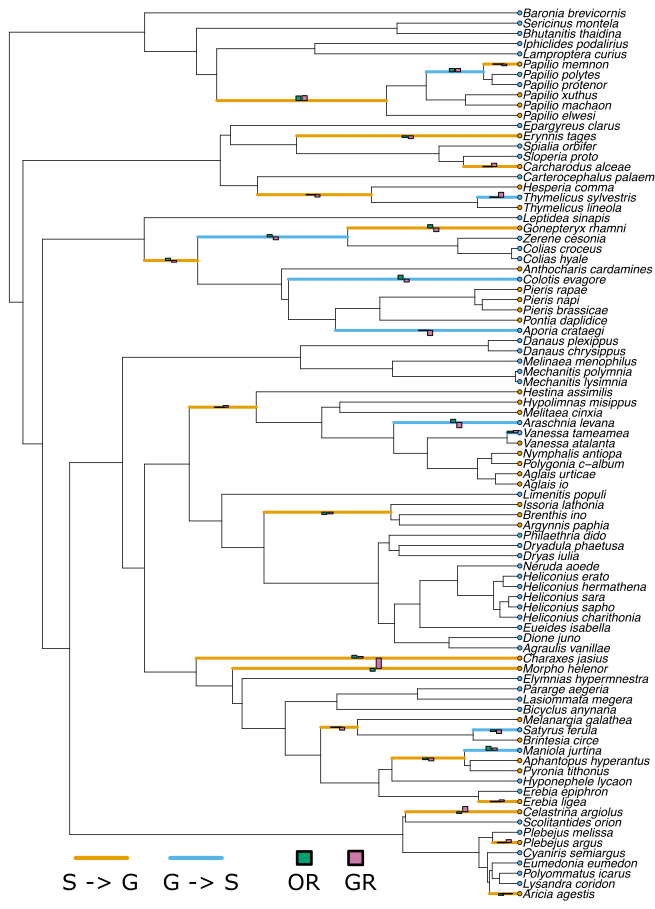
**Figure S4. OR and GR gene family repertoire changes on host plant specialization transition branches.** Specialist to generalist transitions are represented by orange branches, while generalist to specialist are blue branches. OR and GR expansions and contractions are represented by bars above and below branches, respectively. Bar heights are square-root scaled according to the absolute magnitude of net repertoire change to improve visualization of smaller expansion and contraction events.

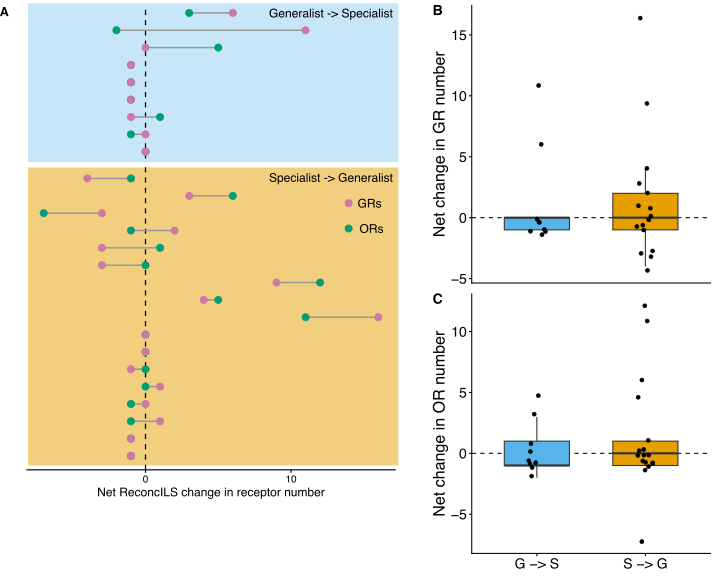

**Figure S5. reconcILS-inferred changes in chemosensory receptor repertoire size associated with transitions in host-plant specialization.** (A) Net changes in olfactory receptor (OR) and gustatory receptor (GR) gene-family size inferred by reconcILS for branches undergoing transitions from generalist to specialist (blue) or specialist to generalist (orange). Each row represents an independent host-specialization transition; points indicate net OR and GR change and connecting lines identify changes occurring on the same branch. Positive values indicate net expansion and negative values indicate net contraction. (B–C) Distribution of net changes for G→S and S→G transitions for GRs (B) and ORs (C), respectively. Net changes did not differ significantly between transition directions for either GRs (ANOVA, *F*₁,₂₄ = 0.019, *P* = 0.891) or ORs (*F*₁,₂₄ = 0.330, P = 0.571). Points represent individual transition branches and dashed lines indicate no net change.
